# 3D neural network-based particle tracking reveals spatial heterogeneity of the cytosol

**DOI:** 10.1101/823286

**Authors:** Grace A. McLaughlin, Erin M. Langdon, John M. Crutchley, Liam J. Holt, M. Gregory Forest, Jay M. Newby, Amy S. Gladfelter

**Affiliations:** Department of Biology, University of North Carolina at Chapel Hill, NC 27599, USA; Institute for Systems Genetics, New York University Langone Health, New York, NY 10016, USA; UNC/NCSU Joint Department of Biomedical Engineering, University of North Carolina at Chapel Hill, NC 27599, USA; Department of Mathematics, University of North Carolina at Chapel Hill, NC 27599, USA; Department of Applied Physical Sciences, University of North Carolina at Chapel Hill, NC 27599, USA; Department of Mathematical and Statistical Sciences, University of Alberta, Edmonton AB T6G 2G1, Canada; Marine Biological Laboratory, Woods Hole, MA 02543, USA

## Abstract

The spatial structure and physical properties of the cytosol are not well understood. Measurements of the material state of the cytosol are challenging due to its spatial and temporal heterogeneity, the lack of truly passive probes, and the need for probes of many sizes to accurately describe the state across scales. Recent development of genetically encoded multimeric nanoparticles (GEMs) has opened up study of the cytosol at the length scales of multiprotein complexes (20-60 nm). Using these probes to spatially resolve diffusivity of a cytoplasmic volume within a cell requires accurate and automated 3D tracking methods. We developed an image analysis pipeline for whole-cell imaging of GEMs in the context of large, multinucleate fungi where there is evidence of functional compartmentalization of the cytosol for both the nuclear division cycle and branching. We apply a neural network to track particles in 3D, generate surface meshes to project data on representations of the cell, and create dynamic visualizations of local diffusivities. Using this pipeline, we have found that there is remarkable variability in the properties of the cytosol both within a single cell and between cells. By analyzing the spatial diffusivity patterns, we saw an enrichment of low diffusivity zones at hyphal tips and near some nuclei. These results show that the physical state of the cytosol varies spatially within a single cell and exhibits significant cell-to-cell variability. Thus, molecular crowding contributes to heterogeneity within individual cells and across populations.

---

The nature of the cytosol has been speculated about since the first glimpses of cells in primitive microscopes, yet remains elusive today. In 1899, EB Wilson expanded on the possible explanations for the mesh-work appearance of the cytosol and described it as an emulsion in which multiple liquids of different chemical and physical properties coexist (1). With electron microscopy, it became clear that the cytosol is crowded and heterogeneous. It is now well appreciated that the cytoskeleton, endomembrane system, and protein translation machinery can generate a crowded landscape in the cytosol. It is not well understood, however, to what degree the motion of macromolecular complexes on the scale of ~10-100 nm is impacted by crowding. Based on concentration alone, it has been predicted that the cytosol should be crowded for macromolecules. However, the effect of crowding on smaller molecules may depend on the spatial arrangement of larger molecules, and some have argued against crowding based on measurements of osmotic pressure (2). Thus, the material state of the cytosol across different length scales is still a poorly characterized factor in cell biology.

Molecular crowding can alter the fluid properties of the cytosol, which in turn influence the thermal, entropic fluctuations of molecular species within, such as Brownian motion of small molecules. Even at the theoretical level, the precise link between crowding and molecular motion is not fully understood. Stochastic molecular motion ultimately directly affects a substantial fraction of the chemical reactions occurring within the cytosol. Indeed, the standard bimolecular reaction rate is proportional to molecular diffusivity, which can be linked directly to fluid properties like viscosity or indirectly modeled through mathematical techniques such as homogenization and stochastic averaging (3). Crowding may therefore have a critical influence on cell function. For example, increased crowding could lower the effective diffusivity for a certain size range of molecules, slowing their reaction rates. Conversely, some reactions may be potentiated by crowding through entropic depletion attraction effects, which favor molecular interactions (4) and molecular condensation (5).

The effect of crowding on particle motion is particularly relevant for large, multinucleate cells, where distinct functional territories emerge within the cytosol. In certain multinucleate fungi, including *Ashbya gossypii*, nuclei divide asynchronously, indicating that proteins that control the division cycle do not diffuse uniformly. Our previous work showed that this asynchrony is in part due to the condensation of RNAs and proteins important for the cell cycle into liquid-like droplets in the vicinity of nuclei (6–9) (see Fig. 1). Multiple studies of liquid-liquid phase separation (5, 10–12) showed that phase separation can be enhanced by crowding, and we speculated that heterogeneity in crowding can influence where biomolecular condensates form.

**Fig. 1.**
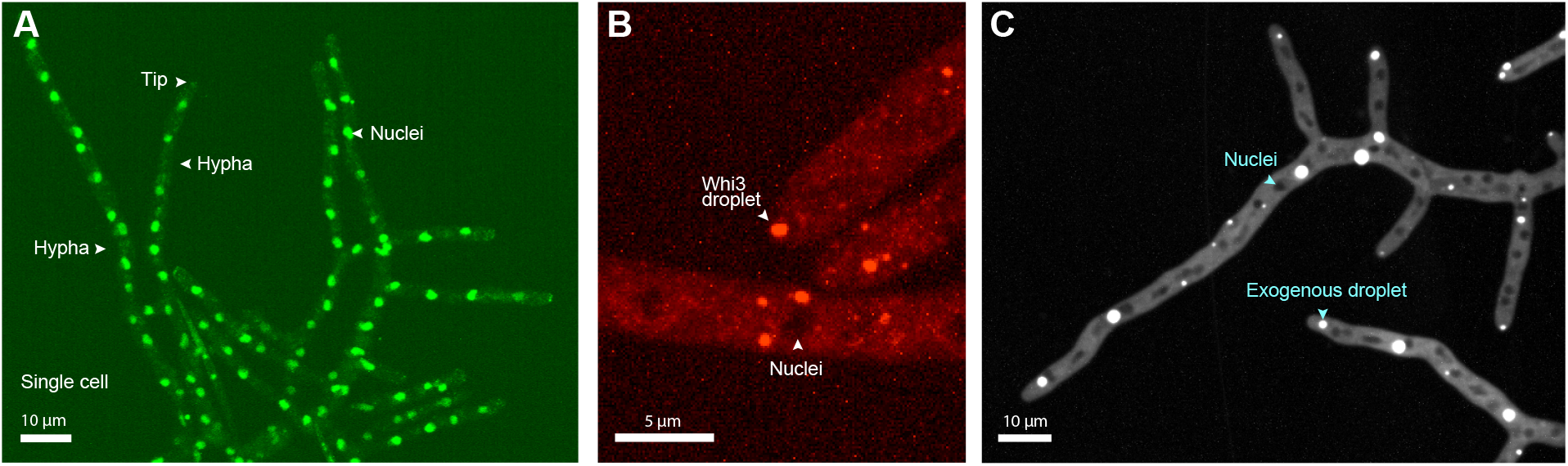
Native and exogenous condensates are non-randomly localized. A. Part of a large, multinucleate *Ashbya* cell. *Ashbya* grows in branched filaments called hyphae, each containing many nuclei residing in a common cytosol. Cells are made up of multiple hyphae and grow exclusively at regions called tips. B. Droplets comprised of endogenously expressed RNA-binding protein Whi3 localize near nuclei and at hyphal tips. C. Exogenous synthetic phase separating proteins are also largely located in the vicinity of nuclei and hyphal tips. Approximately 75% of 113 scored exogenous droplets across 43 hyphae were found to reside within 1μm of nuclei or hyphal tips, measuring from the center of the droplet to the edge of the nuclei or tip.

The best-established methods for evaluating molecular crowding involve particle tracking of ideally passive particles of known sizes that are introduced into cells in a non-invasive manner. They are then imaged rapidly and tracked through time. Based on the observed position-time series tracks, local properties of the medium (e.g., crowding, confinement, viscosity, elasticity, etc.) can be estimated. Applications of particle tracking to measure cytosolic properties in a living cell have been limited due to challenges in creating ideal probes, rapid 3D imaging, and tracking.

Live-cell single particle tracking has previously been applied to track receptors in cell membranes (13, 14). In contrast, we seek to spatially resolve cytosolic properties within whole cell volumes, which requires 3D multiple-particle tracking of inert, genetically expressed probes of known size. The recent development of genetically encoded multimeric nanoparticles (GEMs) has greatly facilitated particle tracking in biological systems. Two types of GEMs have been described, each at a well defined diameter of 20 and 40 nm and icosahedral shape. Introduction of a single gene leads to self-assembly of GEMs in any cell (5). They are an improvement over previous probes such as the *μ*NS system which lacked a stereotypical assembly stoichiometry, size, and shape (15, 16). Recently, GEMs were used to show that changes in ribosome concentration had a substantial impact on molecular crowding and altered diffusivity in the cytosol. The specific 20 and 40 nm size of GEMs was critical for detecting the changes in porosity of the cytosol at the mesoscale length-scale relevant to macromolecular complexes (5). Thus, particle tracking has emerged as a powerful tool for assessing the nature of the cytosol.

There are multiple challenges for 3D particle tracking in live cells using probes based on biological fluorophores. Compared to synthetic nanoparticles, GEMs have lower signal output that, when expressed in *Ashbya* and imaged rapidly in 3D, degrades over time (around five minutes in our case) due to photobleaching, leading to low and temporally decaying signal-to-noise ratio (SNR) conditions. To tackle these challenges, we developed an image processing pipeline that begins with a recently designed neural network particle tracker (17), which is highly automated and was shown to perform well in low SNR conditions. Neural networks have recently made several breakthroughs in microscopy image analysis, including cell tracking (18). Expanding upon the neural network-based particle tracking software, we built an analysis pipeline that constructs a polygonal mesh of the cell surface so that our measurements can be correlated with cellular structures.

Here, we express and track 40 nm GEM particles within the morphologically complex and large cells of the multinucleate fungus *Ashbya* to study the spatial heterogeneity of the cytosol. We find substantial variability in the apparent crowding of the cytosol both within a single cell and between different cells. Furthermore, we find that biomolecular condensates are more likely to form at hyphal tips and in the vicinity of nuclei, regions where we also detected more pronounced crowding. This work provides evidence that the fundamental structure of the cytosol may be an underappreciated source of cell-to-cell variation in populations, which has far-reaching implications for diverse cell processes. This study also provides a technical platform for the spatial analysis of 3D particle tracking data within cells.

## Results

### Native and artificial condensates localize in the vicinity of nuclei and at hyphal tips

How cells control the location of liquid-like compartments formed through phase separation is not well understood. There could be specific molecular determinants that nucleate condensates and/or their positions could be driven by physiochemical parameters such as crowding or pH. The RNA-binding protein Whi3 condenses into liquid-like droplets in the vicinity of nuclei and hyphal tips in *Ashbya* cells to localize transcripts for the cell cycle and cell polarity (6–9) (Fig. 1A,B). We wondered if these localizations are specified by a nucleator of Whi3, or if the physical properties of the cytosol in these areas generally promote phase separation, potentially through molecular crowding. In this case, we would predict that phase separation of other biomolecules would also occur in the same areas where Whi3 droplets are formed.

To test this, we expressed an exogenous synthetic phase separating system made up of multivalent protein interaction domains, coexpressed from a single plasmid developed by Emmanual Levy’s lab [submitted work, personal communication]. Importantly, this system is derived from bacterial proteins and is completely bio-orthogonal to *Ashbya*. Thus, it is highly unlikely that a specific nucleation factor will interact with this synthetic system. We were surprised to see that these exogenous proteins also condensed into droplets in the regions around nuclei and at hyphal tips (Fig. 1C). These droplets span a range of sizes with many much larger than Whi3 droplets, and are positioned such that approximately 75% reside within 1 μm of either a nucleus or hyphal tip. This suggests that the cytosol around nuclei and at tips where Whi3 normally condenses is also able to promote the condensation of artificially engineered condensates. Thus, there may be general features in these areas that promote phase separation. These observations motivated us to examine the physical properties of the cytosol near nuclei and at hyphal tips using particle tracking of GEMs.

### Development of a pipeline for cytosolic particle tracking analysis

To test the hypothesis that there are spatial inhomogeneities in the cytosol, we tracked GEMs throughout the full three-dimensional space of the cytosol in *Ashbya* cells to measure its physical properties. As the tracked motion of nanoparticles can report local crowding, we used the measured diffusivity as a read-out for the state of the cytosol. We also attempted to film smaller 20nm-GEMs, but these were too dim when expressed in these cells, making it impossible to film fast enough (for 3D imaging) to accurately track, so we restricted our focus to the brighter 40nm-GEMs, which we will refer to simply as GEMs throughout the manuscript.

There are multiple challenges for particle tracking in live cells, including acquisition, storage, and automated analysis of 3D video sets. 3D videos can be exceptionally large data sets; a single video can range from ~10-100GB and potentially much larger given the current rapid development in the 3D microscopy space. We implemented our pipeline within Google Cloud to take advantage of their data storage facilities and its full array of high-performance computing services that are designed for high-throughput data processing. The ultimate purpose of the analysis pipeline is extraction of spatiotemporal information of diffusivity in the context of cell architecture (see Fig. 2B).

**Fig. 2.**
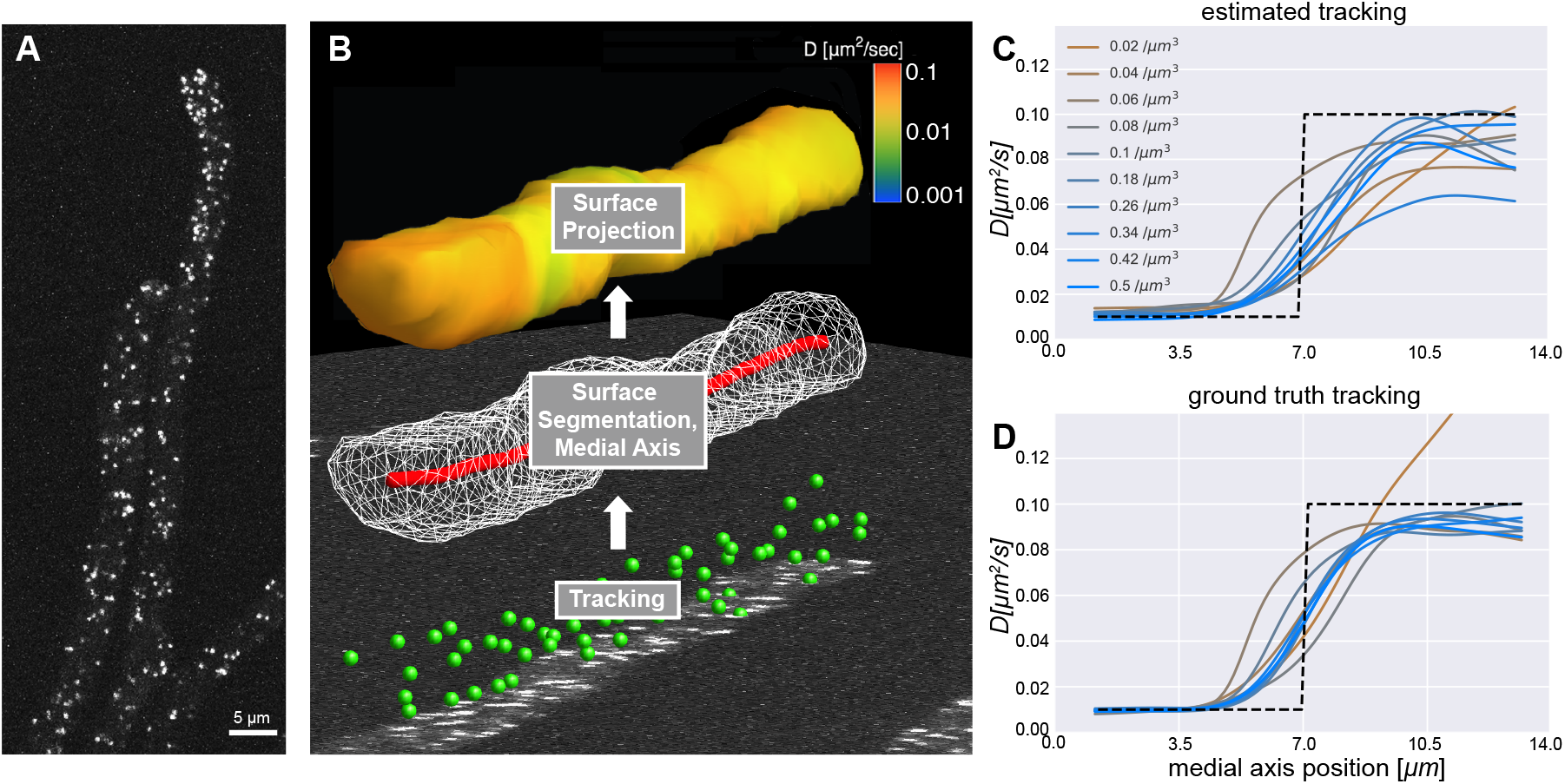
Particle tracking and cellular geometry analysis pipeline. A. A representative max projection of GEMs imaged within *Ashbya*. B. A diagram representing the flow of the analysis pipeline, starting with particle tracking on image data (bottom panel, green spheres), followed by surface segmentation (middle panel, white wireframe of the polygonal surface mesh), and computation of the medial axis (red curve). A representative surface projection of diffusivity is shown for the final step (top panel, heat map). C. Pipeline test 1: Results of analysis of simulations in which hyphae containing GEM-like particles were created to closely mimic our real data. Half the field was comprised of low diffusivity (0.01 μm^2^ s^−1^) particles and half high diffusivity (0.1 μm^2^ s^−1^), shown as a dotted black line. A curve representing the average diffusivity moving along the medial axis is plotted for each simulation. Each curve shows results from simulations with a different density of particles. D. The average diffusivity estimated from ground truth tracks (bypassing the particle tracking stage) that were used to generate the synthetic videos in C.

To spatially analyze diffusivity, we implemented a new processing pipeline built upon a recently developed automatic particle tracking algorithm (17). The neural network tracker employs machine learning methods to accurately localize particle centers from 3D image data (see Fig. 2A). After localization, particle positions are ‘linked’ into tracks. Linking algorithms generally have at least one parameter that fixes the maximum allowable step size between localizations. Because particle mobility varied strongly in space, some regions required small step sizes while others much larger. To adaptively and automatically link a wide range of particle step sizes, we employed a machine-learning method based on the Expectation-Maximization algorithm (19). The localizations were first linked assuming a constant diffusivity, which were then used to estimate the spatially-varying diffusivity. The localizations were then relinked using the spatially-resolved diffusivity. This process was repeated, with each iteration yielding progressively better results. We found that after 12 iterations we achieved sufficient convergence (see accuracy indicated in Figs. 7,8). Additional iterations made virtually no change to the estimates. These methods would ordinarily be computationally expensive to perform on a full set of localizations from a 3D video. By segmenting localizations by hyphae, we were able to link only those that shared the same cytosolic space, as two particles can be close in terms of three dimensional distance but be in different hyphae. The subdivided localizations were also used to compute the cellular geometry for each hypha.

Even with high quality automated tracking software, the end result is a collection of space-time series tracks without the context of surrounding cellular structures. Without this information, we cannot say how a particular particle’s motion relates to the cell structure, or whether it should be combined with or compared to nearby observations. To address this challenge, we used the particle observations to reconstruct the cell surfaces (Fig. 2B) of *Ashbya*, which form highly branched mycelial structures (see Fig. 1). Using the open-source software library CGAL, we used the cell surfaces to compute a medial axis curve that traverses the midline along the interior of the cell surface, which includes the graph structure of branches (Fig. 2B). After obtaining particle localizations, we used the set of 3D points to segment individual hyphae within a given video. We then used the grouped points to construct polygonal meshes of the cell surfaces. The resulting surfaces were then used to compute an approximate medial axis curve, also using the CGAL library (Fig. 2B, middle). The diffusivity estimates are based on averaging the frame-to-frame displacements, or increments, of tracked GEM particles (Fig. 2B, top). The increments are distributed in three dimensions throughout each hypha volume. We take each increment and project it to the nearest medial axis point, the result of which is a kind of histogram, called kernel density estimator (KDE), of the diffusivity along the medial axis.

In addition to the medial axis projections, we projected our estimate to the nearest surface point. This allows us to visualize a heat map of the diffusivity overlaid on the surfaces using the 3D-capable interactive data visualization software DataTank (20) (see Fig. 2B, top, 3A, and Supplemental Videos 1 & 2). While the surface meshes contain more three-dimensional information than the medial axial projections, they also have more uncertainty due to lower sample size. The particle tracking observations are more spread out on the surface than they are on the medial axis. We found both methods to be useful for characterizing the variability in diffusivity over the full dataset. To correlate our diffusivity estimates with the position of nuclei, we projected the nuclei locations to their nearest medial axis point. The medial axis projection also provided a way to analyze diffusivity along the length of the hyphae, allowing us to compare the environment at hyphal tips to the rest of the hyphae.

**Fig. 3.**
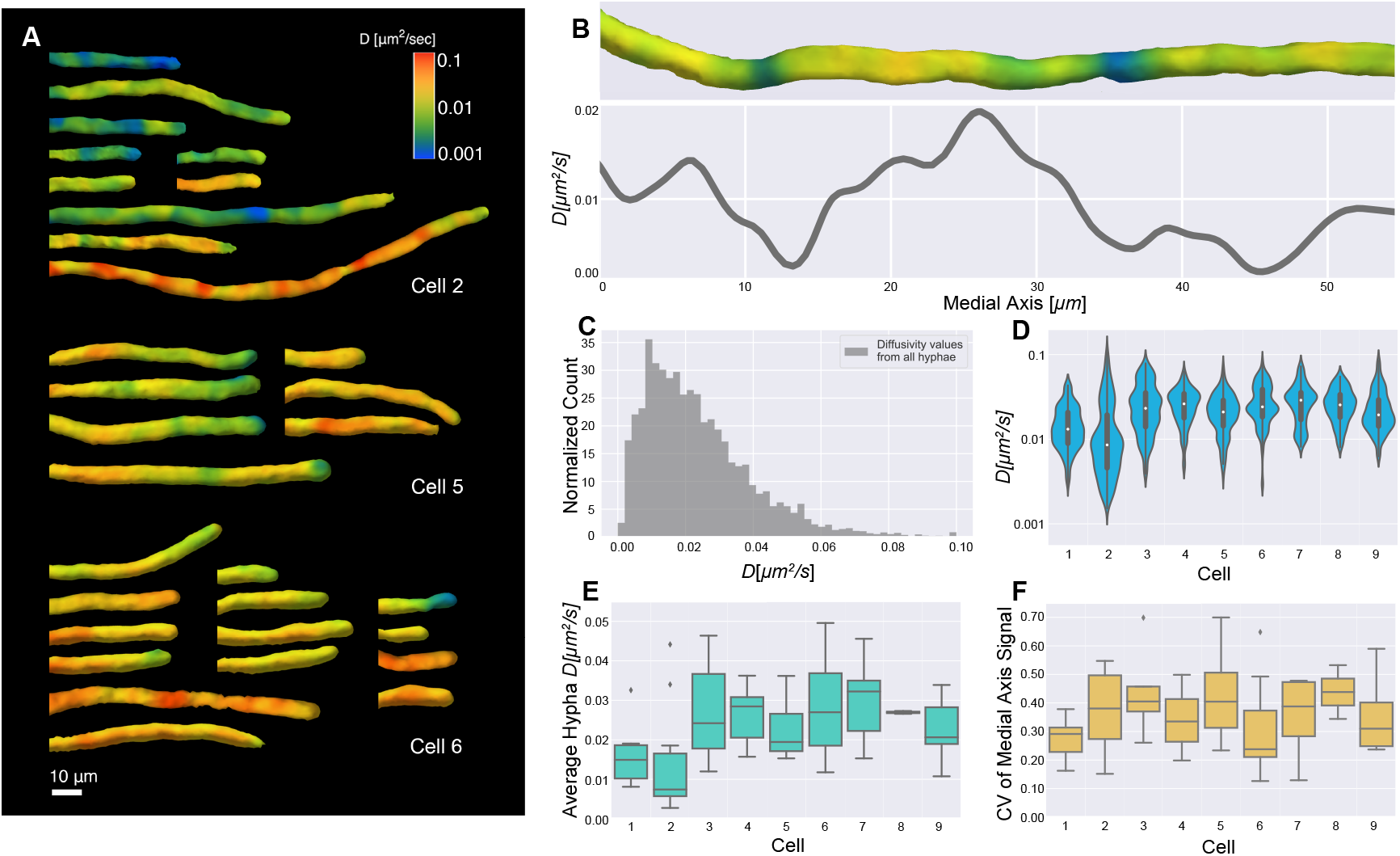
Diffusivity varies within hyphae and between cells. A. Surface diffusivity projections from three cells in the analysis. Note that segmented hyphae were manually aligned for visualization. B. Diffusivity surface projection and corresponding medial axis projection for a representative hypha. C. Histogram of all medial axis diffusivity values from every hypha. 9 cells were imaged with 63 hyphae in total. Average diffusivity is 0.02±0.02 (SD) μm^2^ s^−1^ and n=8558 for medial axis diffusivity values from all hyphae. D. The log-scaled distribution of the medial axis diffusivity values from all hyphae for each cell, indicating overall cell-cell variability. Each cell had 7 hyphae on average, with the number of medial axis diffusivity values per cell ranging from 370-1700. E. Average of each hypha’s medial axis signal, indicating hypha-to-hypha variability of average diffusivity within each cell. F. Coefficient of variation for each hypha’s medial axis signal. Each CV provides a measure of the variability of diffusivity along the medial axis in a given hypha.

#### Testing inference accuracy using synthetic videos

To test the accuracy of the pipeline, we generated videos of simulated data of particles moving in a 14 μm long cylindrical hypha with a 1.7 μm radius (see Fig. 6). The synthetic videos were generated using a custom Python script that simulated particle trajectories and computed the corresponding images that closely approximated the visual characteristics of experimentally-derived videos. Videos were comprised of a time series of 50 z-stacks with an average acquisition time of 0.85 s per stack. Each z-stack had 17 slices with separation of 0.2 μm. Each image slice contained 128×128 pixels with a pixel spacing of 0.11 μm. Three dimensional videos were carefully constructed to closely mimic the video conditions we observed, including noise, 3D particle PSFs, stochastic motion between z-stack slices, and random background intensity within the hypha. All particles moved by simulated Brownian motion with spatially variable diffusivity (consistent with Fickian diffusion). Particle diffusion in each half of the hypha was set to a different value (0.01-0.1 μm^2^ s^−1^).

A set of 10 videos with a range of particle densities were generated to test the accuracy of the medial axis diffusivity estimates. The synthetic videos were fully processed through the particle tracker and the subsequent pipeline (see Supplemental Video 3). Medial axis projections of the test data are shown in Fig. 2C. Error in the absolute measurement of diffusivity was partially due to missing observations and linking errors at the initial particle tracking stage. For comparison, we also used ground-truth particle tracks (bypassing the initial particle tracking step) to isolate the subsequent pipeline error from particle tracking error (see Fig. 2D). Error in the diffusivity estimates using the ground truth tracks was due to sample size and spatial averaging.

Additional synthetic video test sets were generated to assess performance over a range of imaging speeds and diffusivities (see Figs. 7,8) with similar results. Overall, we were able to accurately estimate diffusivities between 0.001-0.2 μm^2^ s^−1^. Lower diffusivities may be possible but were not tested or observed in our experiments. We underestimate high diffusivity values due to time lost imaging through z, but this is a trade-off to enable 3D tracking. We expect the accuracy to improve and better estimates of high diffusivity to emerge in the future as faster 3D imaging techniques are developed.

The measured diffusivities were able to capture the relative change in diffusivity within the synthetic hyphae with sufficient spatial resolution to distinguish each region, which was the primary goal of the pipeline. Importantly, the measurements maintained consistent performance over a wide range of particle densities at the SNR conditions observed in real videos. In summary, these simulations gave us confidence that the pipeline is able to track 3D particles of the intensity and density seen in the data we collected from live cells.

### Diffusivity varies within and between cells

We next analyzed *Ashbya* cells expressing 40nm-GEMs (5) using the pipeline described above. Fig. 3 shows spatially and temporally averaged diffusivity estimates through the first 50 frames, around 40 seconds into imaging, as diffusivity estimates have converged by this time and photobleaching is still negligible. We found that average GEM diffusivity was 0.02±0.02 (SD) μm^2^ s^−1^ (Fig. 3C), slower than previously reported for the same particles in the budding yeast *S. cerevisiae* (5). Our diffusivity is underestimated due to the limitations of 3D imaging rate, but direct comparisons between the species in identical imaging conditions also show that diffusivity in A. gossypii is ~2-fold slower than S. cerevisiae (G.P. Brittingham personal communication). Notably, we found substantial spatial variability in diffusivity, both within individual hyphae and between cells, which are made up of many individual hyphae connected by a common cytosol (Fig. 3A-F).

In Fig. 3A we show three representative montages of cell surface projections of GEMs diffusivity, demonstrating that cells show varying degrees of heterogeneity within hyphae. The diffusivity within single hyphae (e.g., Fig. 3B showing a representative medial axis projection) was observed to range roughly over 0.01-0.1 μm^2^ s^−1^, overall. Given the testing we performed on the pipeline, our measurements likely underestimate the total range of diffusivities somewhat. Nevertheless, it is the relative variability that is of primary interest. In terms of spatial resolution, we were able to detect significant changes in diffusivity over distances as small as 1 μm, even in the medial axis projections which used a 2X longer length scale for KDE averaging compared to the surface projections.

In Fig. 3C we show a histogram of measured medial axis diffusivities, weighted by volume, of the combined dataset. To get a picture of the cell-to-cell variability, the global histogram can be compared to violin plots in Fig. 3D, which show diffusivity distributions grouped by cell. To further quantify the cell-to-cell, intra-cell, and intra-hyphae variability, we computed the average diffusivity per hypha, and the coefficient of variation (CV) of each medial axis signal, defined as the standard deviation divided by the mean (Fig. 3E,F), respectively. Hyphal diffusivity averages varied considerably both within the same cell and between cells. The distributions of CV’s show that some cells (e.g., cell 5) exhibited much more intra-hyphal heterogeneity of diffusivity than others (e.g. cell 1). These analyses indicate that there is substantial spatial variation in crowding as measured by GEM diffusivity both within a single cytosol and between individual cells.

### Diffusivity is correlated with nuclear position and cell cycle

With the observation of spatially heterogeneous diffusivity, we next asked how the variability is organized relative to cellular structures such as nuclei and hyphal tips (see Fig. 4A,D). Videos of cells with both GEMs and a fluorescently tagged component of the Spindle Pole Body, which is embedded in the nuclear envelope, were collected to localize nuclei in different stages of the cell division cycle in relation to diffusivity estimates. As nuclei in *Ashbya* move relatively slowly (21), and all analysis was done on spatially averaged diffusivity estimates through the first 50 frames of GEMs imaging, the nuclei-localized diffusivity values are representative of the local environment around nuclei and unlikely to be impacted by the motion of the neighboring nucleus.

**Fig. 4.**
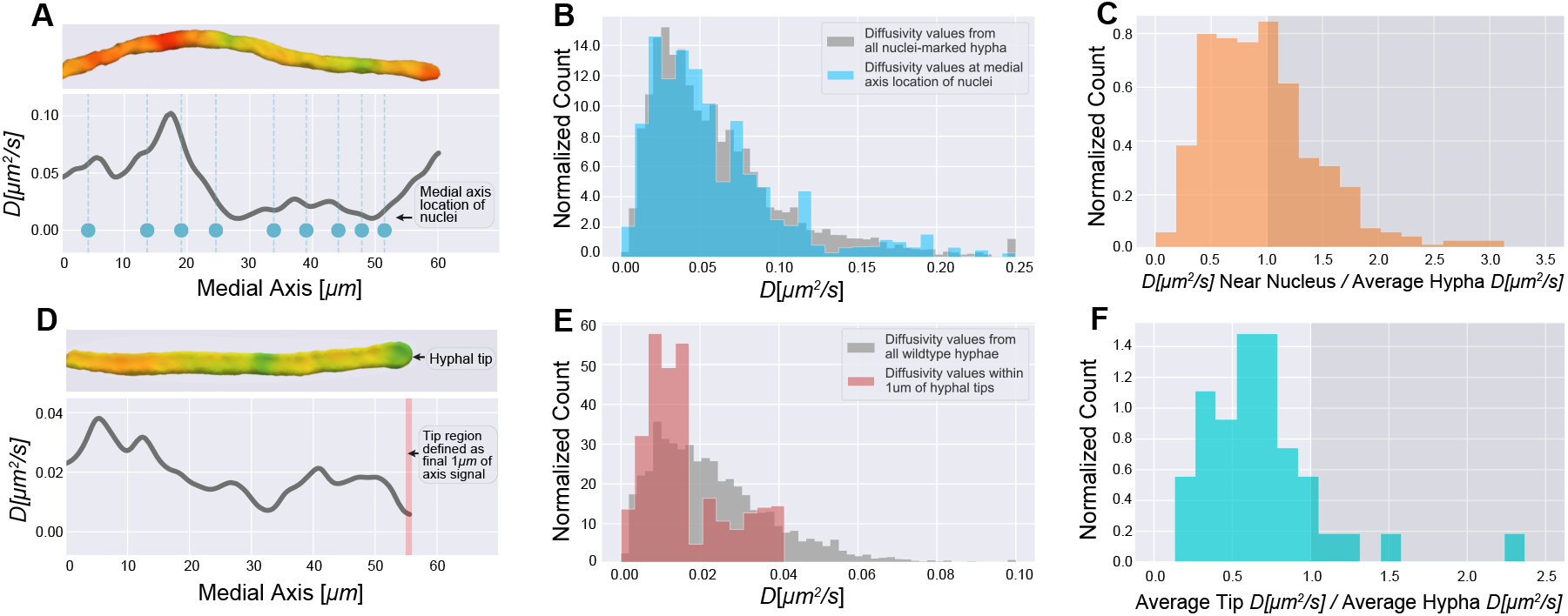
Diffusivity is lower at hyphal tips. A. Single hypha diffusivity surface projection and corresponding medial axis projection. Blue dots show medial axis locations of nuclei within the hypha. Nuclei were imaged first in a single z-stack, followed immediately by a video of GEMs spanning 1-2 minutes in duration. 353 nuclei were localized within 59 total hyphae from 4 cells. B. Diffusivity values from all nuclei-labeled hyphae compared with diffusivity values at medial axis locations of nuclei. Nuclei were localized through fluorescently tagging a component of the Spindle Pole Body. We observed a slightly higher range in estimated diffusivities in this strain compared to wild type. n=9458 for medial axis diffusivity values from all hyphae, and n=353 for medial axis values from nuclei locations. C. For each of the 353 nuclei, we calculated the ratio of diffusivity around nuclei to average hyphal diffusivity. D. Single hypha diffusivity surface projection and corresponding medial axis projection for tip compartment. The red bar highlights the tip region which we have defined to be the last 1um of the medial axis diffusivity curve. For these measurements, 41 hyphal tips were analyzed across 9 cells from the large dataset used for analysis in Fig. 3. Diffusivity values from all hyphae compared with diffusivity values within the tip region. n=8558 for medial axis diffusivity values from all hyphae, and n=226 for medial axis diffusivity values from tip regions. F. For each of the 41 hypha with tips analyzed, we calculated the ratio of average diffusivity in the tip region to average hyphal diffusivity.

Initially, we noticed that some of the low diffusivity regions in the surface projections appeared to be in the vicinity of nuclei. To assess if there was indeed a systematic association between slower than average diffusing GEMs and nuclei, we compared the distribution of GEMs diffusivities in the immediate vicinity of each nucleus to the full spatial distribution of GEMs diffusivities weighted by volume. Any deviation of the former distribution, if statistically significant, would suggest heterogeneity of the physical properties of cytosol around nuclei. Surprisingly, as Fig. 4B shows, both distributions are nearly identical (up to sampling error) however this is comparing to the whole distribution of values rather than comparing regions within a single cell. Therefore, we next compared the ratio of diffusivity in the immediate vicinity of each nucleus to the average diffusivity of the hypha it resided in to make a more local comparison of relative crowding differences. With this local comparison, 59% of these ratios were below 1, suggesting a slight tendency for diffusivity to be locally lower near nuclei. Because the SPB is a reporter of the cell cycle stage of a given nucleus, we also checked if there was any connection between the cell cycle stage and the local diffusivity that might be masked when the data are aggregated for all cell cycle stages. For nuclei in G1 and M, 54% and 53% of the ratios were less than 1, whereas for S/G2 67% were less than one, indicating a cell cycle dependent change in local cytosolic crowding that is most pronounced at mitotic entry. Overall, these results suggest a moderate correlation between nuclei and lower diffusivity compared to the hypha average.

### Hyphal tips are more crowded than other regions of the cell

In *Ashbya*, cell growth occurs exclusively at the hyphal tips, which are known to be sites of enriched actin assembly and liquid-liquid phase separation (see Fig. 1). To quantitatively compare the diffusivity at tips to the overall average, we defined the tip region to be the last 1μm of the medial axis curve (see Fig. 4D), which contains information from the final 2-3μm of the hyphal tip, as medial axis curves did not extend all the way to the surface of tips, but typically ended ~2μm short. Overall, diffusivities at hyphal tips were lower than those measured throughout the full volume of cytosol (see Fig. 4E). Looking within individual hyphae, we observed that a majority of hyphal tips (~90%) had lower diffusivity relative to the average within their hypha (see Fig. 4F). We suspect a small fraction (~10%) of tips have higher diffusivity because they are not actively growing, and the organization of cytosol may depend on the local growth rate.

## Discussion

A major frontier in cell biology is to map the structure of the cytosol to understand how cells may use and react to its different physical states. Although differences in organization can have profound influences on a molecule’s ability to localize to distinct sites in a cell and interact with particular partners or substrates, little is known about how cells actively control the crowding of their cytosol to tune molecular diffusion. Here we find significant variations in the diffusivity of GEM particles within a continuous cytosol and between genetically identical cells indicating that cytosolic organization can be highly heterogeneous. This finding is important because it indicates that a potentially underappreciated source of regulation and noise in biological systems can arise from the physical structure of the cytosol.

As we observed self-assembly of artificial condensates near nuclei and at hyphal tips, our main goal was to correlate cytosolic heterogeneity with the positions of these relevant cellular structures. Compared to their individual hyphal averages, approximately 90% of hyphal tips were found to have lower diffusivity. Tips are known to be sites of actin dynamics, enrichment of ribosomes, and liquid-liquid phase separation which all may contribute to local crowding. Taken together, our results suggest a link between crowding and low diffusivity at hyphal tips and a more modest association between diffusivity at nuclei that is most pronounced in S/G2 phase of the cell cycle. This period is associated with growth and could reflect enhanced ribosome density locally as it is known that the amount of local cytosol per nucleus increases through progression of the nuclear division cycle (22). Why do we not see a more substantial correlation between nuclear areas and low diffusivity zones? It is possible that in this area phase separation is more driven by specific molecules that nucleate condensates or chemical cues that are not detected here. Additionally, there could be a small volume of crowding surrounding nuclei that is a minor fraction of the total local volume of cytosol, making it hard to distinguish by volume averaging estimates. It is also possible that changes in cytosolic properties near nuclei are too small to be detected by our current analysis or are not accessible to the GEMs due to small pore sizes.

This work shows substantial heterogeneity in the motion of 40nm-GEMs in a single cell. What is the source of this variation in motion? The initial characterization of GEMs supported that they are not interacting substantially with native proteins indicating that they should display Brownian motion when the cytosol porosity is larger than 40 nm. We therefore infer that low diffusivity zones are an indication of crowding rather than elastic effects from GEMs interacting with subcellular structures. How do low diffusivity zones arise and persist? We observed significantly lower than average diffusivity within regions as small as 1 μm, and smaller regions may be beyond our ability to resolve without further improvement in the particle tracking methods used here. These regions were relatively stable and often persisted over the full duration of the video (~1-2 minutes). There are several possibilities for what might be causing low diffusivity regions, including crowding by large macromolecules and liquid-like droplets. If untethered, large macromolecules and droplets would diffuse and eventually spread out evenly throughout the cytosol, which would eliminate any heterogeneities. However, for large enough macromolecules, mixing could take a sufficiently long time that crowded regions could persist on the timescale of minutes or longer, consistent with our findings. Moreover, it is possible that some low diffusivity areas are GEMs reporting from within droplets. The interior of RNA liquid droplets has been measured to have substantially higher viscosity than the bulk average we observed within the cytosol (8). Biopolymer networks would also be capable of restricting molecular mobility. Although, it is unclear if such polymeric structures within fungal cells have pore size sufficiently small to affect diffusion of molecules smaller than 50 nm, as the actin and microtubule networks are far less dense than in animal cells.

Substantial improvements in particle tracking are on the horizon, from improved tracking methods, diffusivity estimation, and microscope and camera hardware. A recent study claims to achieve accurate diffusivity estimates and Hurst exponents from short tracks using neural-network-based regression (23), which could allow for less KDE averaging and finer spatial resolution with the same quantity of tracking data. Using light sheet microscopy would reduce photobleaching and allow for longer videos. Currently, imaging speed for 3D microscopy with standard piezoelectric stepping is slower than the framerate of a typical camera, and many groups are working to eliminate this bottleneck (24, 25). Higher acquisition speeds will generally further improve the quality of all tracking inferences, will enable tracking of smaller rapidly diffusing probes, and will allow for acquisition of thicker volumes. These improvements, combined with the engineering of different sized probes with different chemical features, will bring substantial insights into how cells control and use cytosolic organization.

## Materials and Methods

### Plasmid and Strain Construction

Plasmid expressing exogenous phase separating peptides were gifted from E. Levy lab, modified to have a selectable marker for *Ashbya* expression and transformed under standard conditions to generate strain AG893 (26). For nuclear visualization in (Fig. 1A), strain AG275 was used. Whi3-Tomato strains (AG834) were generated by transforming the plasmid AGB 050 into wild-type *Ashbya* strain (AG416). For pPfV-Sapphire GEMs::GEN strains, we transformed the plasmid AGB 910 (pPfV-Sapphire GEMs under control of the ScHIS3 promoter) into *Ashbya* wild type (AG416) and cells expressing Tub4-mCherry (AG270.1) via electroporation and selection on AFM+G418 plates. This generated strains AG837(WT) and AG908(Tub4).

### Cell culture and Microscope setup

*Ashbya* cells AG837(WT) and AG908(Tub4) expressing the 40nm-GEM plasmid were grown under selection of G418 (150μg mL^−1^) in 10mL Ashbya full media (AFM) in a 125mL baffled flask shaking at 30°C for 16 hours. Cultures were transferred to 15 mL conical tubes (VWR) and spun at 300 rpm for 2 minutes. Cells were then washed with 2× Low Fluorescence media, spun again, and placed on a gel-pad embedded in a depression slide comprised of 2× Low Fluorescence media and 1% agarose. Slides were sealed with valap and imaged on a spinning disk confocal microscope (Nikon Ti-Eclipse with a Yokogawa CSU-X spinning disk module and PI P-736.ZR2S triggered piezo stage) using a 100× 1.49 NA oil immersion objective and sCMOS 95% QE camera (Photometrics prime 95B). 3D time lapses of GEMs were acquired using triggered 488 nm lasers at 100% power for 1-2min with a z-stack volume of 3.2 μm (0.2 μm per slice) and 40 ms exposure per image. The average time to image through a volume was 0.92 s, with 200ms on average spent resetting the piezo. For cells expressing Tub4-mCherry, nuclei were imaged prior to GEMs with a single, equivalent z-stack using a 561 nm laser at 40% power with 200 ms exposure. Cells expressing exogenous phase separating peptides were grown and prepared for imaging under the same selection and conditions as cells expressing GEMs. These slides were then imaged using a 60× 1.49 NA oil immersion objective over a z-stack volume of 3.2 μm (0.2 μm per slice) using a 561 nm laser at 100% power with 400 ms exposure. Cells with Whi3-tomato tag were also grown and prepared in the same way, and imaged over a z-stack volume of 3.2 μm using 561 nm laser at 80% power with a 200 ms exposure.

### Figures

Plots were generated using the open-source Python packages, Matplotlib and Seaborn. Images were generated using the open-source application ImageJ and Nikon’s proprietary Elements. Three dimensional surface projections and supplemental videos were generated using the proprietary application DataTank. Adobe Photoshop and Illustrator were used to compile the main figures together.

## SI Appendix

### Tracking analysis pipeline

The GEMs video dataset is comprised of two subsets: WT+GEMs (AG837) and tub4-mCherry+GEMs (AG908). The main purpose of the analysis pipeline is to track GEMs and use the tracks to infer properties of the cytosol, namely, spatiotemporally-varying GEMs concentration and diffusivity. The analysis pipeline is comprised of custom software (written in Python and C++), combined with a number of open source software libraries. In Fig. 5, we show the basic steps in the pipeline.

**Fig. 5.**
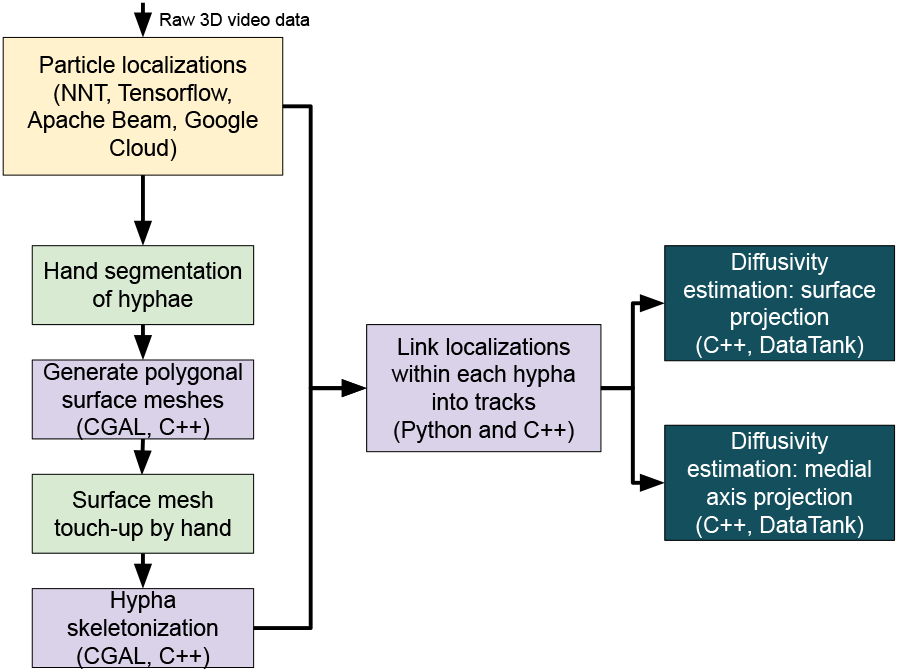
A diagram showing the general organization and steps of the particle tracking analysis pipeline.

The first step is to perform particle tracking on all videos, using the Neural Network Tracker (NNT). The NNT uses map-reduce-style methods (Apache Beam) together with cloud computing resources from Google Cloud (Dataflow) to batch process our large datasets and large file sizes.

The second stage of the pipeline is to compute an accurate reconstruction of the cell surface and a coordinate system representing the cell geometry. Because of the branched morphology of the cells, multiple branches often lay side by side, making it difficult to group GEMs localization belonging to a specific cell. In order to compute the GEMs concentration, we also need to estimate the local volume of cell interior, which requires some means of determining whether a given point is inside or outside the cell and which hypha it belongs to.

Once we fully construct cell geometries for each hypha segment, we use it to group GEMs localizations by hypha segment. We then link the grouped GEMs localizations through time into paths or tracks. This ensures that a given particle path will not jump from one hypha segment to another.

We then use kernel density estimation and exponential time averaging, to compute spatiotemporal estimates of diffusivity. We use two geometric projections for these estimates: a medial axis projection and a surface projection. In each, observations of particle movement are grouped together based on their location. For example, in the medial axis projection, we group observations by their closest point on the medial axis. The medial axis projection assumes that the cytosol is approximately radially symmetric around a given medial axis point and that its properties change only along the length of each hypha. The surface projection assumes that cytosolic properties might vary depending on the radial direction but not on the radial distance from the surface. The medial axis curves have fewer points than the surfaces, which means the medial axis projection results in reduced sampling error compared to the surface projection because, on average, more observations are grouped together compared to the surface projections.

#### Hypha geometry estimation

After the videos have been tracked, we use the tracking data to generate a surface mesh and the medial axis curve (including branch points) extending down the length of each hypha. All of the particle localizations (particle positions), are grouped within each video. The resulting point set is used to generate a polygonal surface mesh of each hypha. Because it is exceedingly difficult to automatically segment each hypha (particularly when two hyphae lay alongside one another) hand segmentation and hand touch-ups of the surfaces meshes is performed. Only the non-overlapping portions of hyphae were included, with all including or being near to a hyphal tip. These are the only steps in the pipeline that require interactive processing, the remaining steps are fully automated.

The surface meshes are initially constructed using tools from the Computational Geometry Algorithms Library (CGAL). First, a 3D Delaunay triangulation is computed for each hypha segment, which results in a convex solid comprised of many connected tetrahedra. The cell surface is then computed using the CGAL 3D Alpha Shapes package. The resulting surface mesh is usually quite rough and ill-conditioned for the downstream steps of the pipeline. The CGAL Triangulated Surface Mesh Simplification package is used to resample and smooth the surface mesh. In some cases, topological imperfections cannot be removed automatically (a known problem in constructing surface meshes from random point collections). We typically see 1-2 small spots on approximately one out of every ten hypha segments that require hand corrections. We repair these surfaces meshes using Meshmixer.

The medial axis for each hypha segment is computed using the CGAL Triangulated Surface Mesh Skeletonization package (27). This package requires a well-formatted surface mesh, free of holes and topological imperfections. An approximation of the medial axis is obtained, a connected graph of points, which extends along the center of each hypha, connected at branch points. A secondary product of the mesh skeletonization procedure is a mapping from a given medial axis point to surface points. Each medial axis point is connected to zero or more surface points such that each surface points is connected to exactly one medial axis point.

This data structure has several uses. It can be used to construct a method for determining if a given point is inside or outside a given hypha segment. We also use this mapping to project a given point within a hypha segment to the closest medial axis point. This allows us to generate estimates of diffusivity along each hypha medial axis. We also obtain estimates of concentration and diffusivity on the cell surfaces by mapping a given interior point to the nearest surface point.

#### Using polygonal surface mesh and medial axis to sort points inside a given hypha segment

We use the surface mesh and the medial axis curve to efficiently determine if a given point is inside or outside. While this is trivial to compute for closed convex regions, hypha segments are almost never convex. They are, however, locally convex. Let the medial axis be given by *M* = Γ × *E*, where Γ = {**x**_*k*_} is the set of nodes and *E* = {e_*j*_} is the set of edges linking each node. As described above, we also have a mapping from medial axis points to local surrounding surface points. Let *S*_*k*_ = {**x**_*m*_} be the set of surface points connected to the medial axis point **x**_*k*_.

Given an input point **x** to test, the method proceeds as follows. First, the closest medial axis point **x**_*A*_ to the test point **x** is computed. This step can be done quickly by brute force as there are a relatively small number of medial axis points. Second, we search the set of surface points connected to **x**_*A*_ for the closest local surface point **x**_*S*_. These are given by

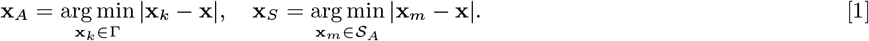

Then, we say that the test point x is inside the hypha segment if

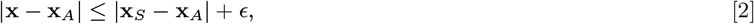

where *ϵ* > 0 is a small parameter that pads the estimate so that points that are very close to the surface are not mistakenly excluded. The parameter *ϵ* can be thought of as the uncertainty in the estimated surface position. In pixel coordinates, we use *ϵ* = 1, which works out to be approximately 100 nm.

#### Kernel-density estimation with exponential time averaging

Spatiotemporal estimates of diffusivity are computed using a kernel density estimator. This can be thought of as a ‘smooth histogram’. Let the set of particle positions at time *t* be given by {*X*_*n,t*_}, for 0 ≤ *n* ≤ *N*_*t*_, where *N*_*t*_ is the total number of particles at time *t*. A Gaussian centered at each observation with a predetermined length scale parameter *σ* (~6μm for surface projections and ~12 μm for medial axis projections) are summed together. For the first stage in computing estimates (prior to time averaging), we define

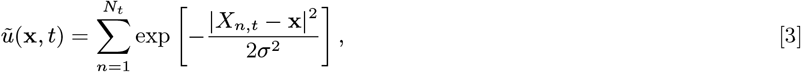

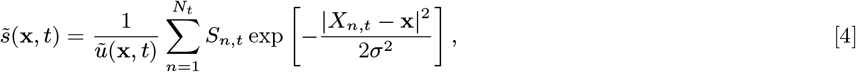

where *S*_*n,t*_ is the contribution to the estimate from a single observation, centered at *X*_*n,t*_. For diffusivity, the standard maximum likelihood estimator is given by setting 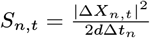, where 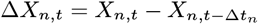 is a particle displacement (or increment) and Δ*t*_*n*_ is the time step between observations within the track, which is an integer multiple of the average inter-stack time interval of the video (the linker can skip over missing observations).

The next step is to apply exponential time averaging to the result. For a given function *f* : ℝ^*d*^ × ℝ_+_ → ℝ, let *f*_*j*_ = *f* (**x**, *t*_*j*_), for 1 ≤ *j* ≤ *M*, where *t*_*M*_ = *T* is the max time. Then, the final time-averaged estimates are defined by

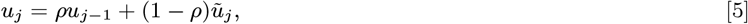

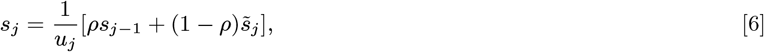

where *ρ* ∈ (0, 1). We can relate *ρ* to a physical timescale with 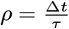, where Δ*t* is the average inter-stack time interval of the video and *τ* is a parameter that determines the time-scale of averaging. We use timescales of *τ* = 64s and 192s for exponential averaging of surface projections and medial axis projections, respectively.

#### Radius estimation and filtering large particles

We observed a small number (compared to GEMs) of fluorescent particles with larger radii than GEMs within some hyphae. Their origin could not be determined. The distribution of these larger particles was random, though possibly elevated somewhat near tips. They moved randomly, diffusively or possibly subdiffusively, with mobility substantially lower than GEMs. To reduce their potential influence on our measurements of heterogeneous diffusivity of GEMS, we estimated the PSF radius and peak intensity of all particle localizations by regression of the local surrounding image patch to a Gaussian profile.

The filter was more accurate when applied to tracks instead of individual localizations. Averaging the PSF radius and peak intensity over all localizations within a track mitigated the influence of noise. We applied basic linking (assuming constant diffusivity for filtering) to the localizations to obtain a set of tracks specifically for filtering. The tracks used in primary analysis were computed during a later step. After linking, we averaged the radius over each track to get the average PSF radius (*r*) and the peak intensity (*I*_peak_). We filtered those tracks that had at least four observations and satisfied at least of one of the following two conditions

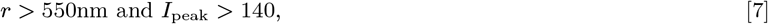

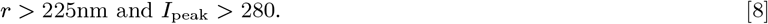

These criteria were chosen heuristically to achieve a balance between filtering as many large particles as possible while minimizing the number of true GEMs mistakenly filtered out. Due to this trade off, some of the larger particles were tracked. However, their contribution to the overall observed heterogeneity was minimal.

Within the wildtype set, over half of the hyphae had no larger particles visible, and 90% of the hyphae had <5% of their localizations coming from the larger particles. All hyphae in the analysis had <10% of their total localizations coming from large particles. Hyphal tips were only included in the analysis if they did not have large particles within last 3μm of the hypha. The larger particles were infrequent in the Tub4-Cherry set, accounting for <1% of localizations.

### Pipeline testing with stochastic particle simulations and synthetic videos

To test the particle tracking pipeline, we generated synthetic videos of particle diffusing in a cylindrical domain with reflecting boundaries. The cylindrical domain was used to mimic a small section of a hypha, approximately 14 μm long and 3.4 μm in diameter (see Fig. 6). The diffusivity was changed at the midpoint of the hypha so that the left half had a lower diffusivity than the right half. Five video sets, each containing 10 videos, were generated with different diffusivity values for the left and right half of the cylinder. We varied the total number of diffusing particles and the SNR for each video within a given video set. Videos where generated with Gaussian noise, slowly varying background intensity (mimicking nonspecific fluorescence within the hypha), and a small ~3 pixel radius Gaussian PSF for each particle (see SI Video 3).

**Fig. 6.**
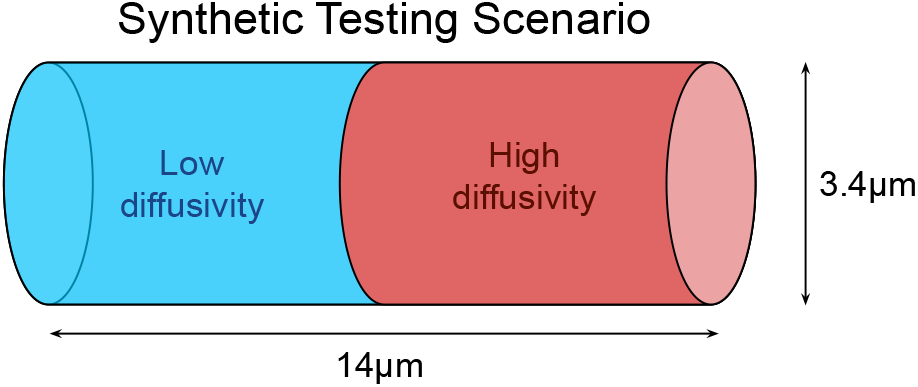
Diagram illustrating the cylindrical domain used to generate synthetic testing videos. The cylinder represents a small segment of a hypha where there is a sharp jump in diffusivity. The purpose of the test is to quantitatively measure the ability of the pipeline to simultaneously measure low and high diffusivity colocalized within a hypha.

**Fig. 7.**
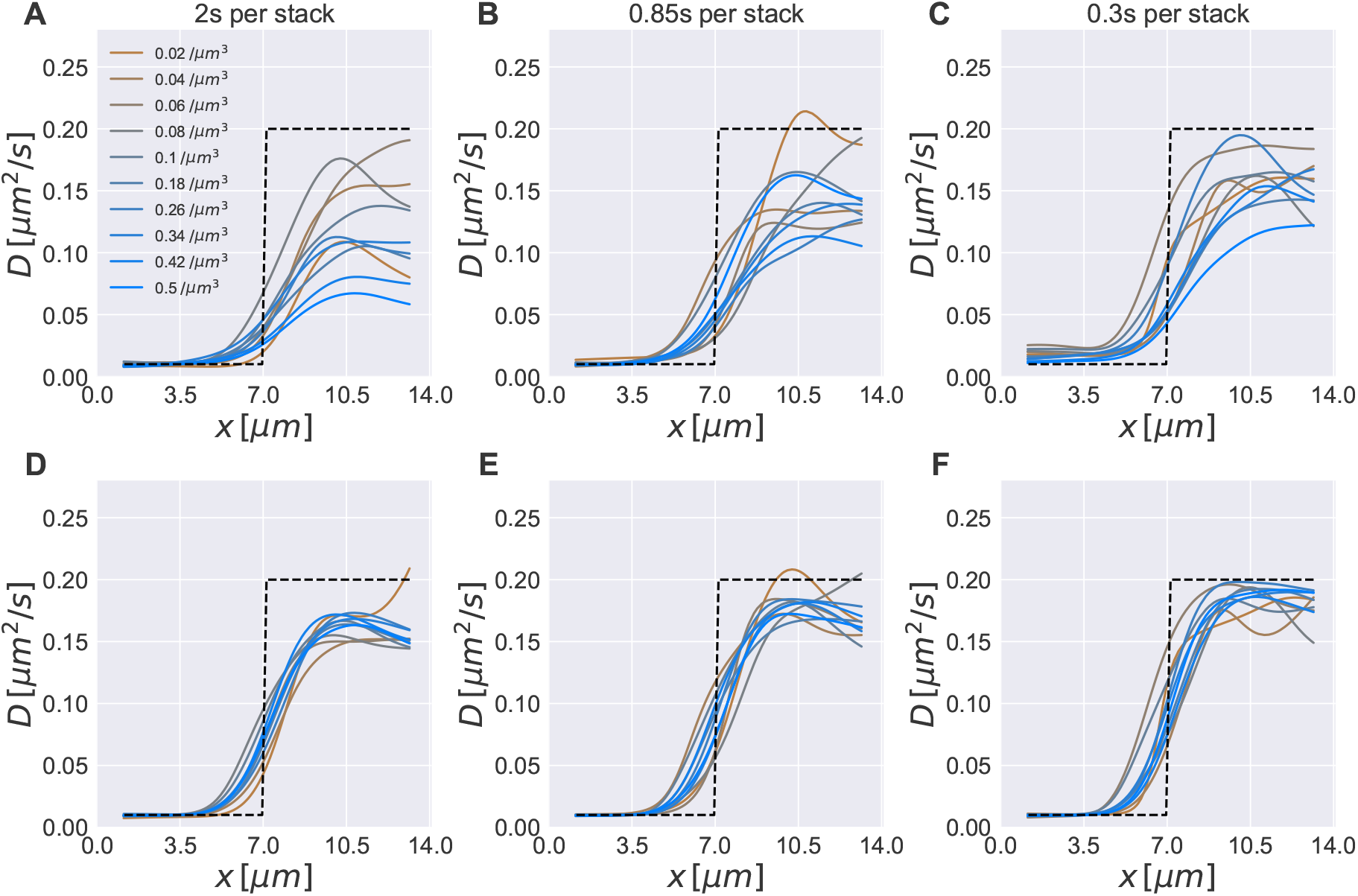
Synthetic testing result show accuracy across a range of imaging speeds. A-C. Diffusivity estimate from particle tracks obtained by processing synthetic videos through the full pipeline. D-F. Diffusivity estimate obtained from processing ground-truth particle tracks (that were used to generate the synthetic videos) through the analysis pipeline. For reference, the GEMs videos were imaged ∼0.85s per stack, and GEMs particle density varied around 0.05 - 0.15 μm^−3^.

#### Stochastic particle simulations

Particle simulations of Brownian motion with spatially varying diffusivity (under the Fickian interpretation) were performed using a time stepping method. We assumed independent particle motion so that there were no interactions between particles. Let *X*_*t*_ ∈ ℝ^3^ be the position of a track at time *t*. The simulation scheme is given by

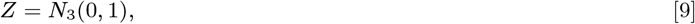

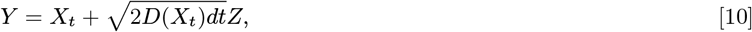

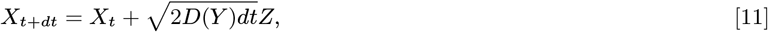

where *N*_3_(0, 1) ∈ ℝ^3^ is a vector of numerically-generated, independent, normal random variables with mean zero and unit variance. The simulation time step was *dt* = Δ*t*/16 (i.e., there were 16 simulation time steps between each z-stack of the simulated video). Reflection at cylinder boundaries was done using the method of ballistic reflection, which preserves detailed balance.

#### Synthetic testing results

To examine the effect imaging speed, given by the average time to image each z-stack, we generated test videos at 2, 0.85, and 0.3 seconds per stack. The GEMs videos were on average around 0.85 seconds per stack. In these tests, the low diffusivity was 0.01 μm^2^ s^−1^ and the high diffusivity was 0.2 μm^2^ s^−1^, which was at the high end of our measurements for intracellular diffusion of GEMs.

We assess the performance of the pipeline (see Fig. 7) by comparing the medial axis diffusivity estimate generated by the analysis pipeline to the ground truth values (dashed line) used to generate the videos. Three video sets are shown at 2, 0.85, and 0.3 s per stack. GEMs videos were imaged at ~0.85 s per stack. The top row (Fig. 7A,B,C) show results from processing the synthetic videos through the full pipeline, while the bottom (Fig. 7D,E,F) row shows result of processing the ground truth particle tracks through the pipeline. The latter is intended to show the estimates obtained with perfect particle tracking for comparison. For each testing set, we accurately estimated the low end diffusivity (left half) and underestimated the high end diffusivity (right half). Faster imaging speed improved the accuracy of high diffusivity, while accuracy at low diffusivity was slightly reduced for the fastest imaging speed (0.3s per stack) due to simulated localization error.

To ensure that we can simultaneously measure very small differences and very large differences, automatically, with the same linker, we varied the range of diffusivities in an additional test set. In Fig. 8, we show a 10× reduction of diffusivities, compared to Fig. 7.

#### Boundary effects on particle tracking

The tube-like hyphal geometry restricts 3D Brownian motion to a quasi-1D closed domain. Using a 3D diffusivity maximum likelihood estimator, the apparent diffusivity from purely 1D Brownian motion would be 4× lower than unbounded 3D Brownian motion. We were able to reduce this potential effect by ignoring the z-position in our diffusivity estimates, which reduced the maximum possible boundary effect from a 4× reduction to a 2× reduction. Boundary effects should begin to affect diffusivity estimates (reducing the estimated diffusivity by up to 2× its actual value) once the average width of a hypha is about the same distance as a typical GEMs displacement during one video frame. In the synthetic testing, we observed ~10% reduction in estimated diffusivity when the true diffusivity was 0.2 μm^2^ s^−1^. At hypha tips, there is another boundary restricting motion, and we observed an additional underestimation effect at high diffusivities (~15% at 0.2 μm^2^ s^−1^). As expected, this effect was negligible for lower diffusivities.

### Surface projection videos

Three videos were generated to illustrate the spatio-temporal surface projection estimates. SI Video 1: Max projection of GEMs in an *Ashbya* cell. 3D GEM localizations are visualized as green spheres above the max projection, and surface projected diffusivity estimates are shown above those. SI Video 2: Max projection, localizations, and surface projected diffusivity estimates for a single hypha. Lower diffusivity at the hyphal tip as well as heterogeneity of diffusivity within the hypha can be seen. SI Video 3: An example of a testing video as described in Fig. 6. Max projection of simulated Brownian motion shown, with localizations as green spheres, and surface projected diffusivity estimates above.

**Fig. 8.**
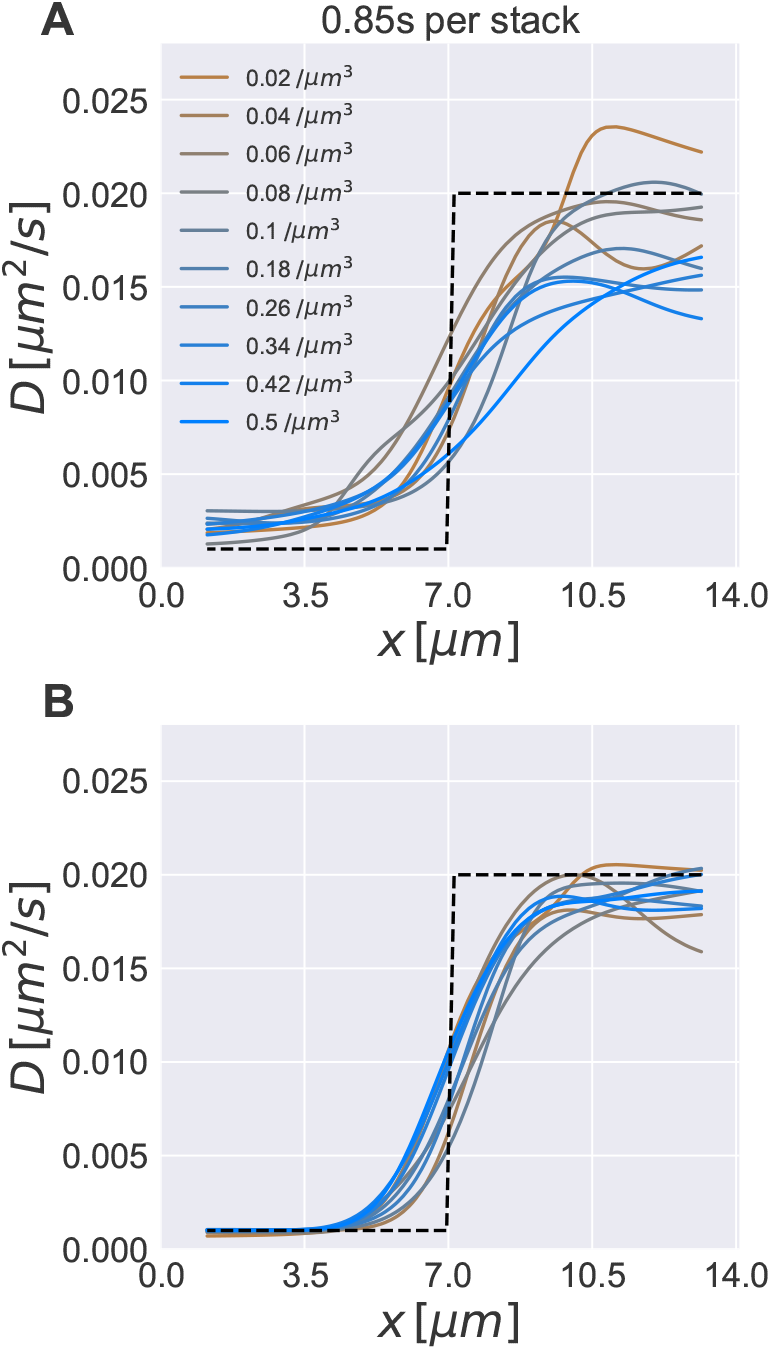
Synthetic testing results show small diffusivity differences can be accurately resolved. A. Estimated diffusivity from processing synthetic videos through the full analysis pipeline. B. Estimated diffusivity from processing ground-truth tracks through the analysis pipeline.

## ACKNOWLEDGMENTS

We would like to thank the 2016 Physiology course at the Marine Biological Laboratory in Woods Hole, MA, Gregory Brittingham, and Marcus Roper for initial experiments and perspectives on pipeline. We thank David Adalsteinsson for help with DataTank software and many conversations about image analysis on large datasets. We thank Emmanual Levy (Weizmann Institute) for providing plasmids encoding synthetic phase separating peptides. This work was supported by Google Cloud, the National Science Foundation (NSF), the National Institutes of Health (NIH), and the Natural Sciences and Engineering Research Council of Canada (NSERC). ASG was supported by the NSF (RoLs: 1840273) and the NIH (R01GM081506). JMN was supported by the NSERC (RGPIN-2019-06435, RGPAS-2019-00014, DGECR-2019-00321) and the NSF (DMS-171474). MGF was supported by the NSF (DMS-1816630, DMS-1664645). LJH was supported by the NIH (R01GM132447).

